# Kinship governs fertility with pre-and post-zygotic mechanisms mediated by match of methylation patterns

**DOI:** 10.1101/135327

**Authors:** M. Herbert

## Abstract

It has been shown that kinship normally determines fertility in humans and animals. At smaller populations the curve of fertility against kinship rises steeply as kinship rises until inbreeding is reached. In large and progressively larger populations, fertility slowly falls. At an equilibrium point fertility matches replacement. Away from that, fertility changes by one of four published time courses. Earlier we demonstrated damped oscillation in a population of captive fruit flies. I hypothesized that since fertility changes so rapidly, DNA base mutations could not be the cause. The mechanism must be epigenetic, the control of genes, and gave varying doses of a methylating cocktail to the same flies. The fly population changed in a complex way, which only happens when there are pre-zygotic and post-zygotic processes in effect together.

**One Sentence Summary:** There is evidence for an epigenetic effect – with both pre-zygotic and post-zygotic components – of kinship on fertility of flies (*Drosophila melanogaster)* induced by a methylating cocktail.

## Main Text

In 1967 Robin Fox published *Kinship and Marriage* (1) in which he examined traditional societies and found that virtually all had stringent rules about who could marry whom. These traditions have been largely abandoned in our globalizing, urbanizing world, but one would suspect that there was a selective advantage to them. More recently Fox has pointed this out in his chapter “Marry in or Die Out” (2). In 2007 Richard Sibly and others published a paper (3) that examined more than a thousand serial field counts of animals and found that population size, the inverse of average kinship, was regularly related to population growth, which is a function of fertility; larger populations were less fertile. Other studies find the same relationship in humans (4, 5, 6). The question arises as to what mechanism could be at work.

We published an account (7) of the time course of a population of captive fruit flies, and saw damped oscillation with a characteristic rapid rise and slow fall, which could be modeled with a computer program employing a post-zygotic mechanism penalizing any virtual couple’s fertility as their kinship diminished. The oscillations of the test population were so rapid that it was clear that awaiting changes in DNA sequence simply could not account for them. The mechanism had to be epigenetic – that is related to some control mechanism of the relevant genes, not the genes themselves. Of course egg and sperm unite to form a zygote; anything that reduces the chance of the sperm arriving or the formation of a zygote is pre-zygotic infertility while anything that intervenes to lower the chances of the zygote undergoing growth and development to form a fully fertile adult is post-zygotic. Since the damped oscillations could be accounted for fully by a post-zygotic process, I thought I might be able to change the fertility of the flies by making an epigenetic change.

Folic acid tends to increase the rate of methyation of DNA, and there is a published cocktail of a number of dietary supplements calculated to enhance the process (8). Since the ingredients are all readily available this seemed like the easiest place to start. Of course given the number of epigenetic processes known, there was no assurance that this was the right track, merely the convenient one. I mixed the cocktail – it proved to be a suspension rather than a simple solution – and tested it on some flies. In my hands, any concentration over a 40% dilution was lethal, so that seemed to be about the place to start and gradually work downward. I simply continued with the same population previously published.

I shall proceed to explain why we should not be surprised that such a mechanism exists, then to show the Sibly curve and compare it with some computer simulation of populations, give some patterns the model can show and compare them with real populations. We shall look at the experimental results and see what they indicate about the (relative) infertility. Finally I shall make some suggestions as to what directions work might go in the future.

Starting from first principles, every animal must have a niche. Niches change. Animals change. A species awaits the appearance of a new niche and since niches close there is selective pressure to occupy new ones.

Consider two animals each occupying a low value niche when a new and rich niche opens. Over generations selection will move them toward the new opportunity so that neither is quite optimized for its original niche. One animal becomes better optimized for the new niche first and monopolizes it so that the other species loses out and returns to its less favorable niche. Thus selection is a race. Now consider a case where the original niches are not that bad. Selection moves each species to a compromise. But one species undergoes speciation, breaks into two species, one of which exploits its original niche and the other optimizes for the new one. The tardy species loses the opportunity. What’s worse, while it is displaced into a compromised condition another species may undergo speciation and claim that old niche as well as its own. The slowly speciating form might go extinct even though its original niche never changed. So speciation is a race.

But rapid speciation comes at a cost. In the absence of a general consensus on how long speciation takes, we shall assume two thousand generations. Imagine an animal in a valley that passes a copy of one chromosome to two different offspring. We will call one chromosome and all its descendants f_m_ and the other line f_n_. For clarity we will assume for now that there is no recombination and no genetic drift; each chromosome passes one and only one copy on in the population. Now f_m_’s carrier hops across the valley, and a glacier descends splitting the valley for two thousand generations. The glacier melts. Now f_m_ may be brought back so its animal can meet f_n_’s and attempt to have offspring. But they cannot have fertile offspring; it has been too long. This is standard allopatric speciation. Next instead of a glacier, assume the population remains constant at 1,000, recalling that no chromosomes enter or leave. If the population mates at random, it will again take roughly two thousand generations for f_m_ and f_n_ to get together. All other pairs of chromosomes are similarly distant from each other or more distant, so the whole population dies. Eliminating some of the chromosomes by genetic drift only raises the number of individuals the population can sustain; the logic remains unchanged. Admitting recombination does not change the logic either.

Selection will tend not to tolerate extinction of a highly successful life form so there is great selective pressure, over the long run, to put in a fix. We can see the fix in action in this curve drawn from Sibly (3). We shall draw it like this, so as not to commit ourselves on numbers we don’t really know: *(fig 1)*.

**Fig. 1.**
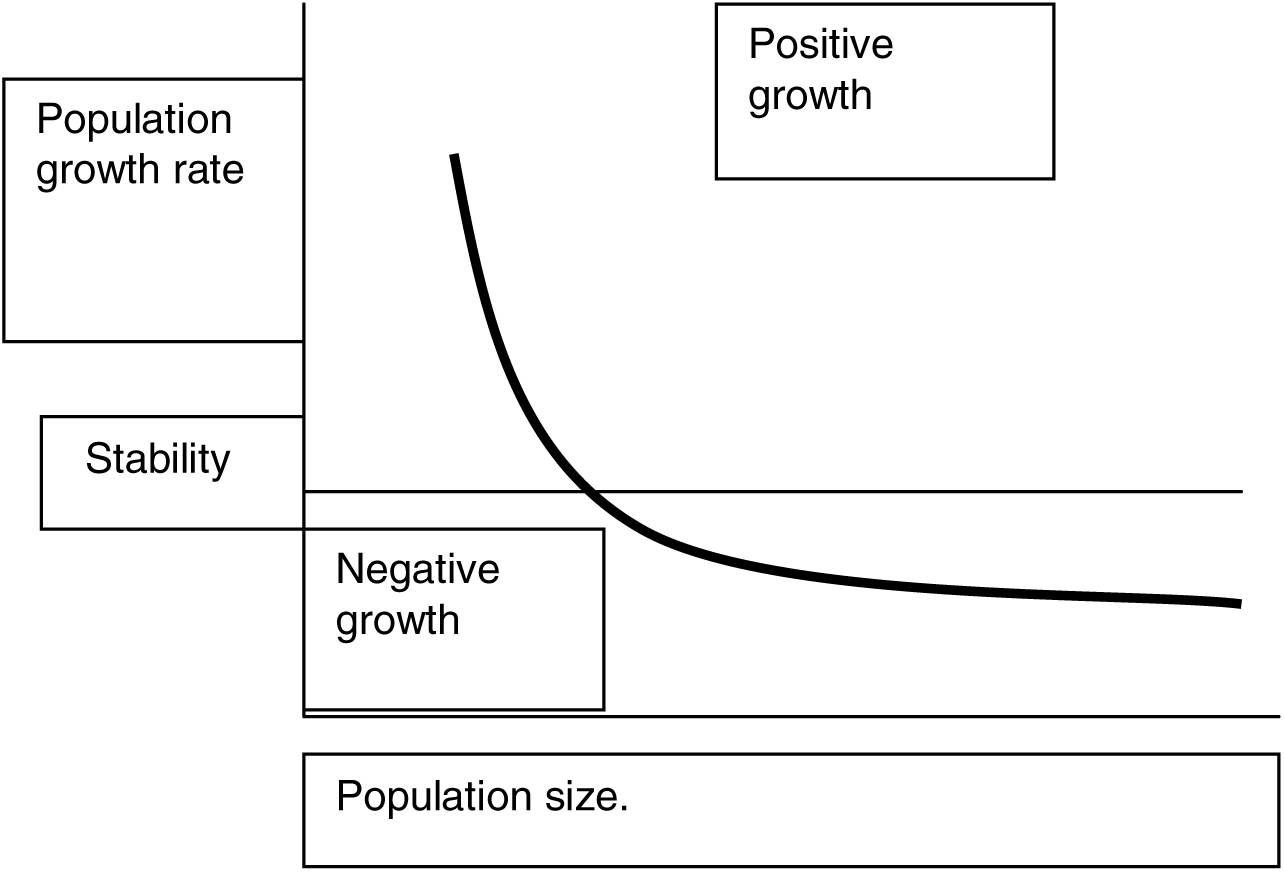
Abstracted from Sibly (3).The vertical axis, ordinate, is the population growth rate increasing upward. The horizontal axis, abscissa, is the population size increasing to the right.

This summarizes or at least abstracts from over 1,000 serial field counts of animals. The same curve can be seen among humans in Iceland (4) both for children and grandchildren and in Denmark(5,6) using distance between birthplaces of couples as a surrogate for kinship. That study uses this “marital radius” as the abscissa, which of course follows the square root of the area and population size, the inverse of kinship; correcting for that, the curves are the same. Crucially, the pair led by Laboraiu looked at the issue of choice.

Once they took town size and marital radius into account, there was neither effect of income nor education of the couple on the number of children they had. This is a biological process, neither economic, social nor voluntary. That is hard enough to grasp, and it turns out to be even harder to bear in mind at all times.

Whether a population remains at rest or changes, and the time course by which it may change, can vary. I cannot offer a grand theory of what conditions produce what patterns.

But I can offer a few patterns exhibited by a computer model. In the first I set up a virtual population subject to pre-zygotic and post-zygotic fertility reduction (parameters laboriously tweaked) more severe with decreasing kinship, ran the population for a thousand generations, put it to bed, got it back up and continued another two hundred generations. The graph is the time course of those two hundred generations, each vertical bar a generation and the height being the total number of offspring the population generated each generation (after which offspring were eliminated if need were to reduce the population size to the maximum that had been selected at the outset, 700. ) *(fig. 2)* to deal with as we look for other patterns.

**Fig. 2.**
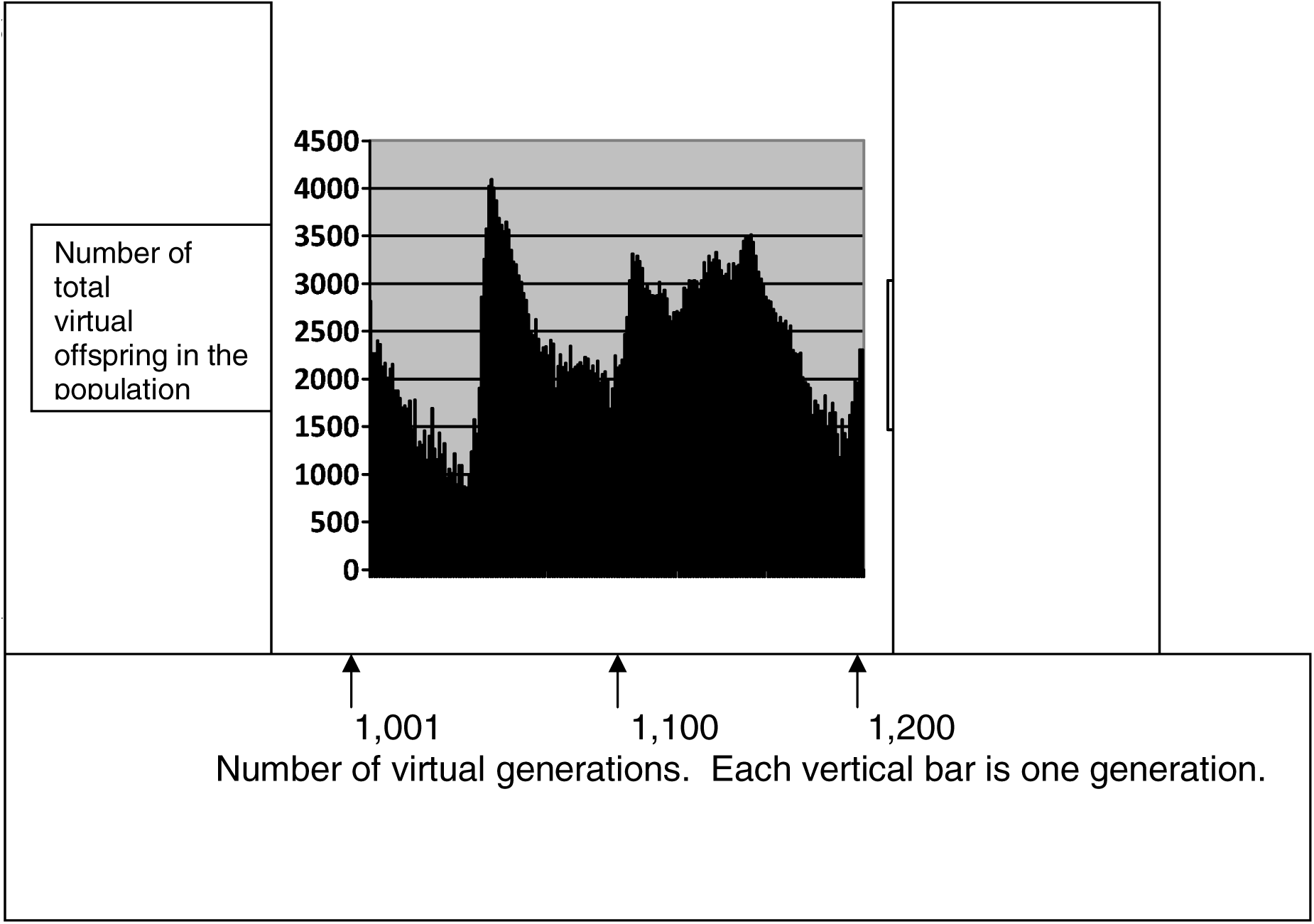
Baseline of a computer simulation of a population regulated by pre-zygotic infertility and post-zygotic infertility both increasing with decreasing kinship. The population was initially run for 1,000 generations with maximum size of 700. The final 700 were then run for 200 more generations shown as 1,001 through 1,200. Each vertical bar is a single generation. The vertical axis is the number of offspring for the entire population in that generation. This is the baseline and is very quite noisy even after having a significant time in which to stabilize.

In the wild, real animals seem to do better. If we were to look at the record of mouse counts in Australia, where they monitor wild mice in order to follow plagues of mice, (9) most of the time the counts are low and stable.

A second pattern is damped oscillation. *(fig. 3)*

**Fig. 3.**
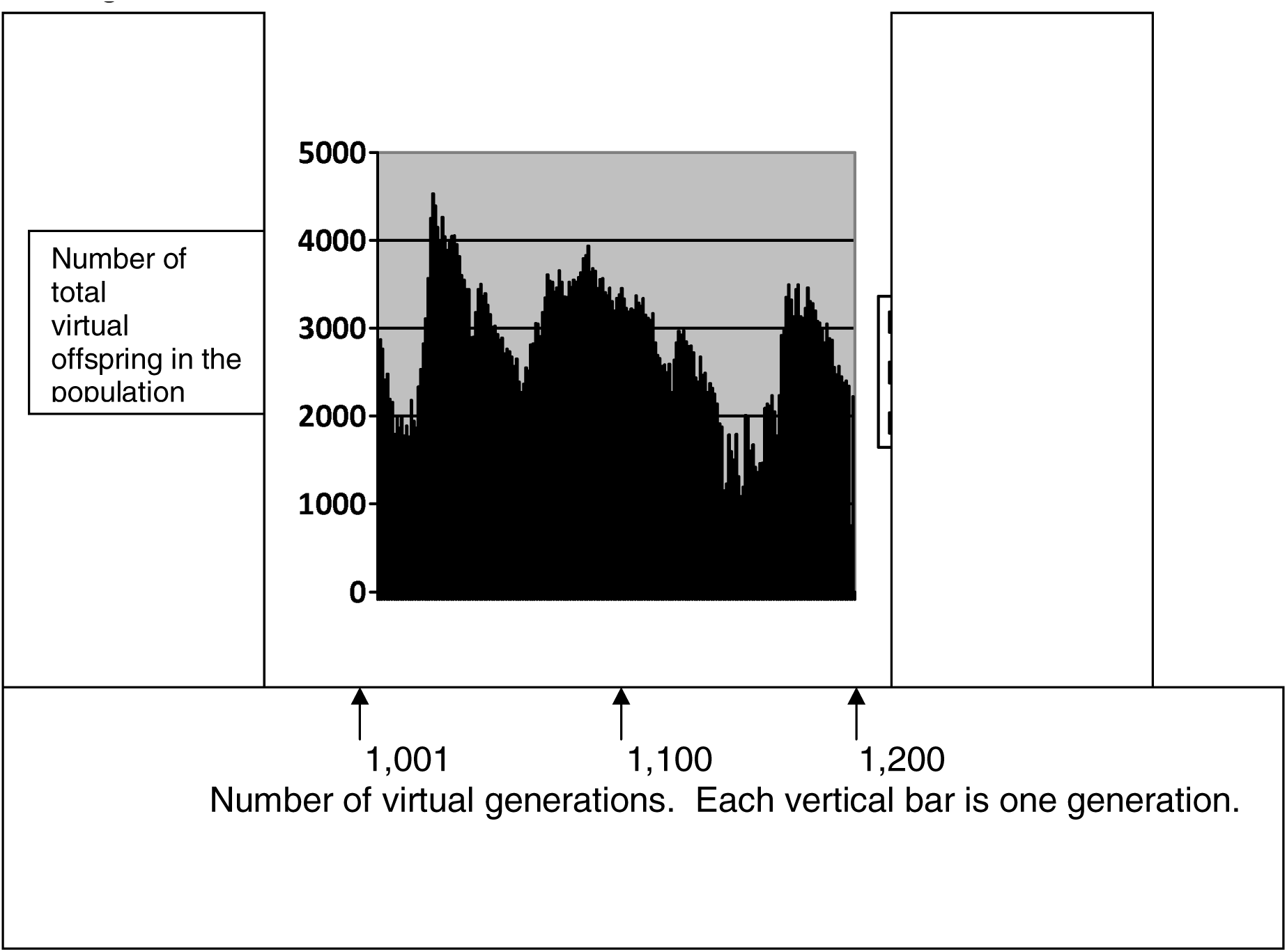
Computer simulation of a run like the one in figure 2, except starting with a smaller population, 200 instead of 700. The vertical axis is the number of offspring for the entire population in that generation. The horizontal axis numbers the generations as 1,001 through 1,200. The pattern approximates a damped oscillation.

This curve was obtained with a run in which only 200 of the saved population were brought out and run; all other conditions were identical with the baseline, and like the baseline the population survived all 200 generations of the test. As with the baseline, the time course is very noisy, but one can make a case for there being three cycles of damped oscillation. Arguably the cycles not only decline but have a characteristic rapid rise and slow fall. Damped oscillation has been demonstrated by computer model by me using only post-zygotic infertility (7), in the laboratory by me using fruit flies(7), and in the wild among European voles (10). I don’t think we quite know enough to make the claim, but I would only faintly object if someone were to declare that the Sibly curve already implies a point of equilibrium such as the mice and rabbits found and suggests oscillation with a rapid rise and slow decline; predicting the damped nature of the oscillation would seem more difficult.

A third curve consists of two peaks ending in population collapse. *(fig.4)* The population died out after 102 generations. I discount the trifling rally of the last few generations; we know the baseline is unstable. Conditions were identical with those of the baseline run except that maximum population was permitted to rise to 3,000.

**Fig. 4.**
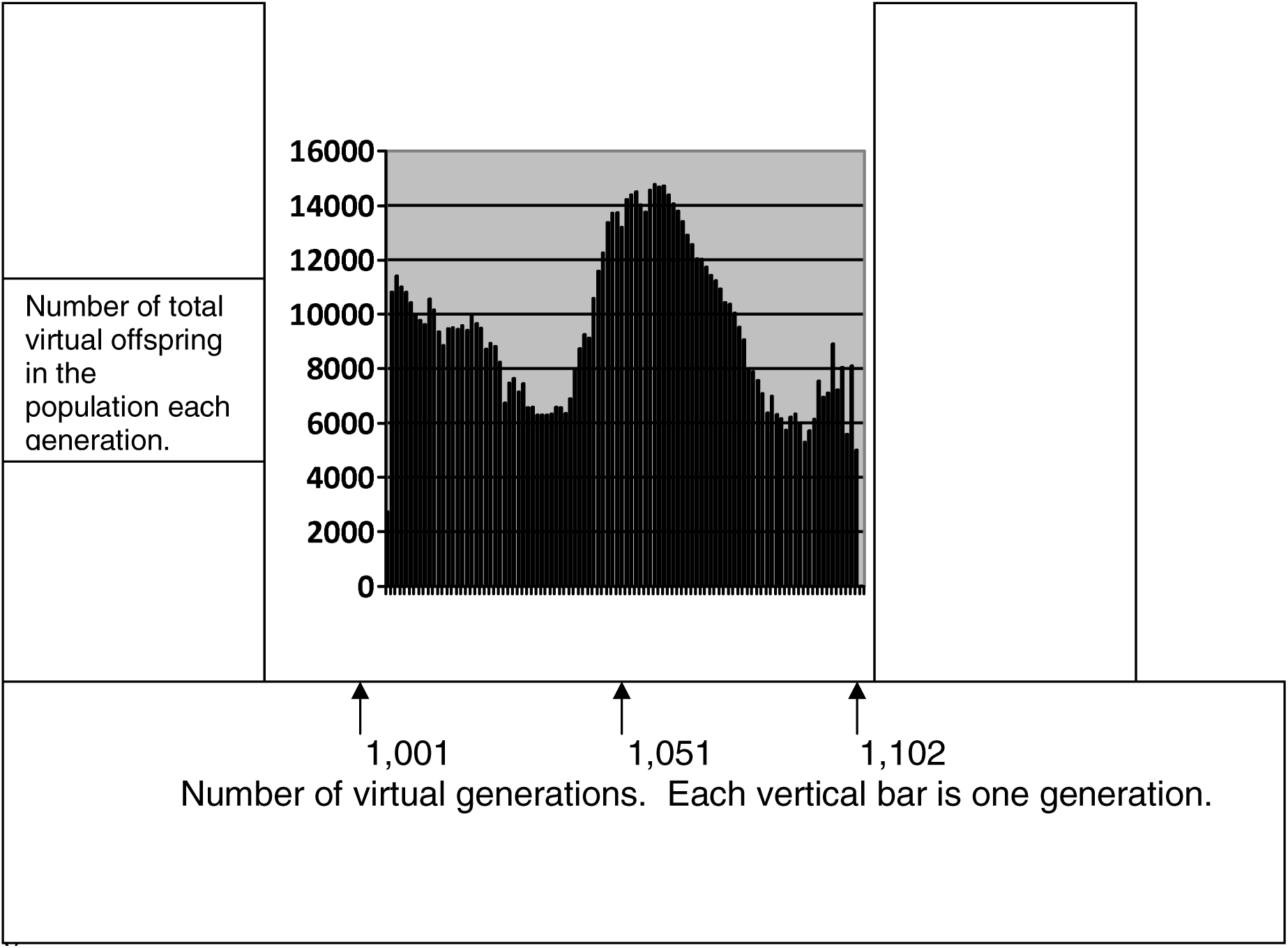

This is complex behavior that I have not been able to elicit neither with a pure post-zygotic mechanism nor a pure pre-zygotic mechanism. It is not something I would expect to occur had I only the Sibly curve to look at. Yet it happens in real life. In the Australian mouse counts(5), the generally low stable populations are occasionally punctuated by spectacular rises in the numbers of mice. These typically follow a drought. That, of course, makes good sense; the population has been shifted to the left on the Sibly curve. The mice are never without food, nor are the available predators sufficient to stuff enough mice into their mouths to tame the plagues. It might be disease, but there is no evidence for that. Twice in the published record there is this same profile: two peaks, the second higher than the first. If you look up the original article, have a bit of caution; it may look like there are three double peak plagues, but one of them is the overlap of counts from two different stations.

This same double peak occurs among humans. Jarred Diamond published an article about Native American farmers living in Long House Valley in the Southwest somewhat isolated in their desert fastness surviving for centuries as a distinct population (11). By locating every cook fire location in every dwelling in the valley and doing C-14 dating on bits of charcoal he was able to produce an annual census. By looking at tree ring widths, he was able to calculate how fast the trees were growing. The two correlated very well so he reached the quite reasonable conclusion that the population in the valley was being governed by the weather. I wish to cast no shadow on Diamond’s excellent work. Indeed I posted his graph inside my front door for years, telling myself, “Inherent in that curve is the rule that governs love, society, human history and natural history; until you understand it well enough to reproduce it you understand absolutely nothing of any consequence.”

Forgive my enthusiasm (I was younger then) but what I noticed was that in early days there were stepwise increases in the population size; people were moving in and doing so multiple households at a time. There were no obvious stepwise declines, so emigration cannot have occurred, nor significant war, famine nor plague. In fact during early years people were moving in at a time when the weather, under the climate hypothesis, would have meant they starved, but they flourished just as well as during other years of that era. The people were excellent farmers; no European has ever been able to farm in that valley. They knew years ahead, and they were cultivating the trees. That may have involved nothing more complicated than taking wood uniformly throughout their groves rather than proceeding by clear cutting at the edges. At all events, it was a population of humans with no outside influence save the original immigrations and following only the logic of its fertility mechanism.

And that mechanism produced the two peak curve ending in population collapse. It has been suggested that all empires fall when they reach an age of 250 years (12). Living in a nation that started around 1776, I do hope that’s wrong. But if we look at the Long House Valley experience, when the population was about 500 they were accepting another 200 immigrants. That might indeed be enough people to form a nucleus of ruling elite of a substantial population. If that is typical for the beginning of an empire, then its downfall would match the Long House Valley collapse. History is not my field, but I find it hard to think global collapse awaits us only nine years hence.

A fourth curve was produced by letting the maximum population rise to 4,000, *(fig. 5)*, but otherwise keeping conditions identical with the baseline run and the run with two peaks.

**Fig. 5.**
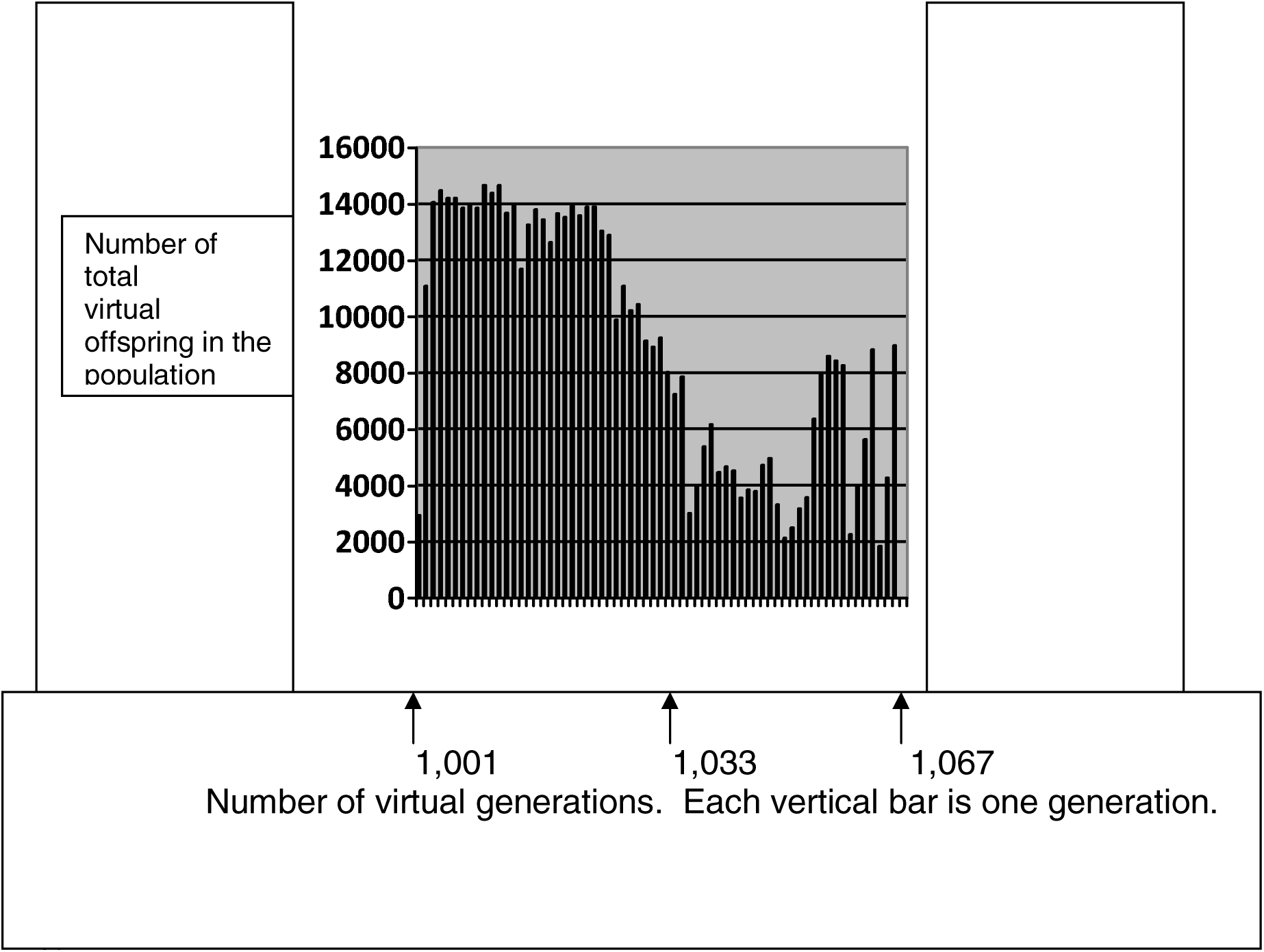
Computer simulation of a run like the one in figure 2, starting with the whole population of 700 but raising the maximum population size to 4,000. The vertical axis is the number of offspring for the entire population in that generation. The horizontal axis numbers the generations as 1,001 through 1,102. The pattern shows one major peak and a small rally at the end.

It died out after 67 generations. Ignoring the abortive final rally, the pattern looks like fast up, slow down, tends to level off and then collapse. Given the difference in resolution, this is consistent with the four biggest mouse plagues in the Australia experience (9), with lab work done by Calhoun (13), and matches quite well with a computer model, which summates a number of counts of outbreaks of leaf cutting insects in Canada (14). An enthusiast might say, “It’s obvious from the Sibly curve; if the population goes too far out to the right it will go extinct before it drops to the rest point, and at the end as the rest point is approached fertility should improve a bit.” I can only say that the computer program has burnt me many times, and I caution against drawing conclusions before the fact.

The experimental model was the same cage and flies that had previously demonstrated a post-zygotic pattern of fertility declining with population size^7^. I followed the population for more than two years to confirm the baseline. *(fig. 6)*

**Fig. 6.**
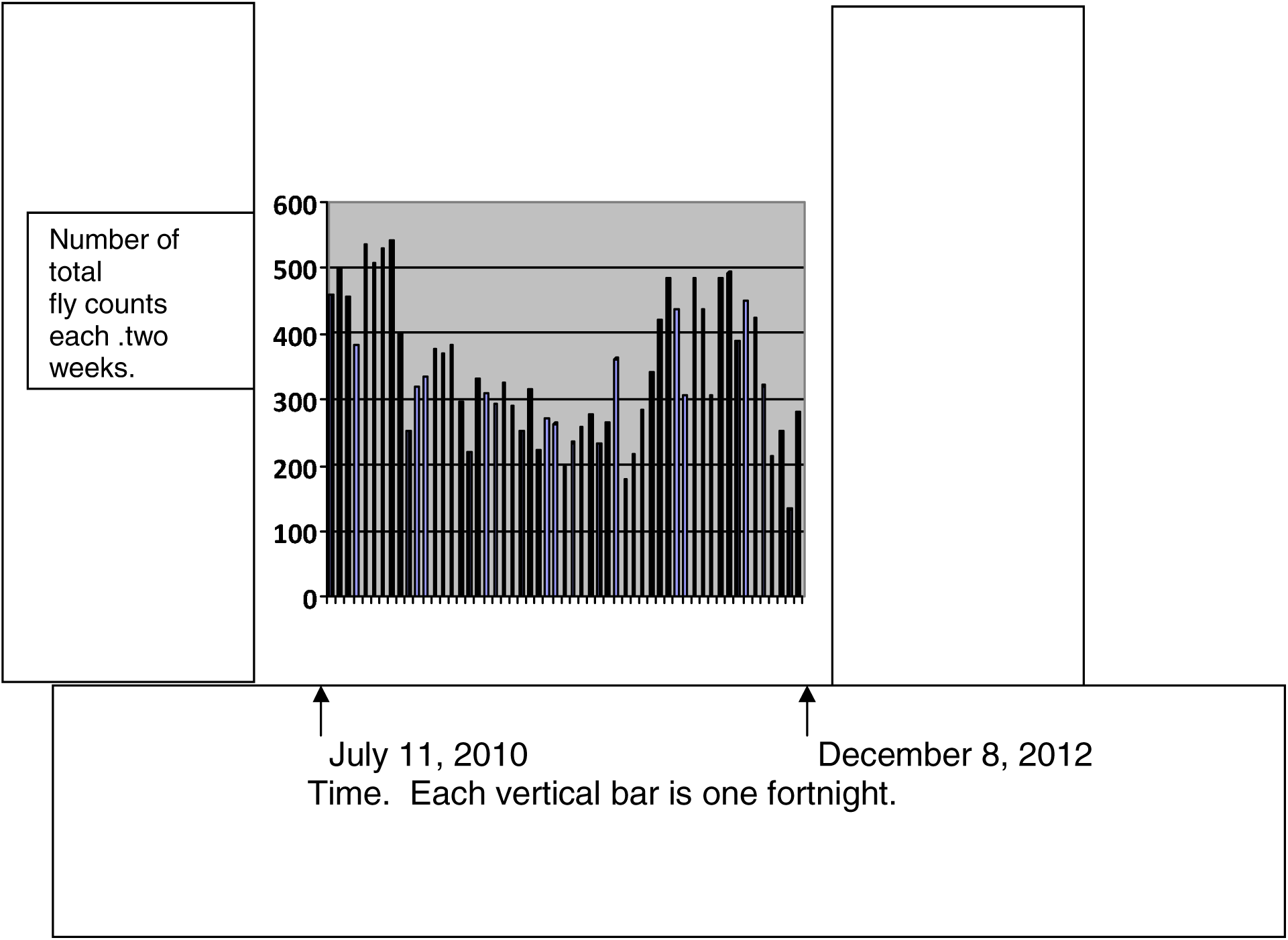
Thirty month baseline of daily counts of a captive fruit fly population through two windows accumulated at two week intervals. Early on there is a rough damped oscillation although this seems to break up in the final year.

Each vertical bar is the total number of daily counts in two windows in the cage summated over a two week interval.

As in the original data, we ascertain a pattern of damped equilibrium for most of the time, although this is breaking up at the end.

Because the fertility of the flies changes too rapidly for DNA mutation to be the cause I was convinced that an epigenetic mechanism was the cause and methylation was the easiest place to start, but I had no idea what dose level might change the fertility of the flies. So I started with a sub-lethal dose and worked downward. *(table 1)*

**Table 1.**

| Dose | Begin | End |
| --- | --- | --- |
| Baseline | July 11, 2010 | December 8, 2012 |
| 40 % | December 9, 2012 | January 5, 2013 |
| 30 % | January 6, 2013 | April 29, 2013 |
| 20 % | April 30, 2013 | July 23, 2013 |
| 16 % | July 24, 2013 | November 13, 2013 |
| 13 % | November 14, 2013 | March 1, 2014 |
| 11 % | March 2, 2014 | June 23, 2014 |
| 9 % | June 24, 2014 | October 3, 2014 |
| 7.5 % | October 4, 2014 | January 6, 2015 |
| 6 % | January 7, 2015 | April 28, 2015 |
| 5% | April 29, 2015 | August 19, 2015 |
| 4% | August 20, 2015 | November 12, 2015 |

Daily counts in two windows continued and the results pooled over 2 week intervals were again examined. *(fig 7)*

**Fig. 7.**
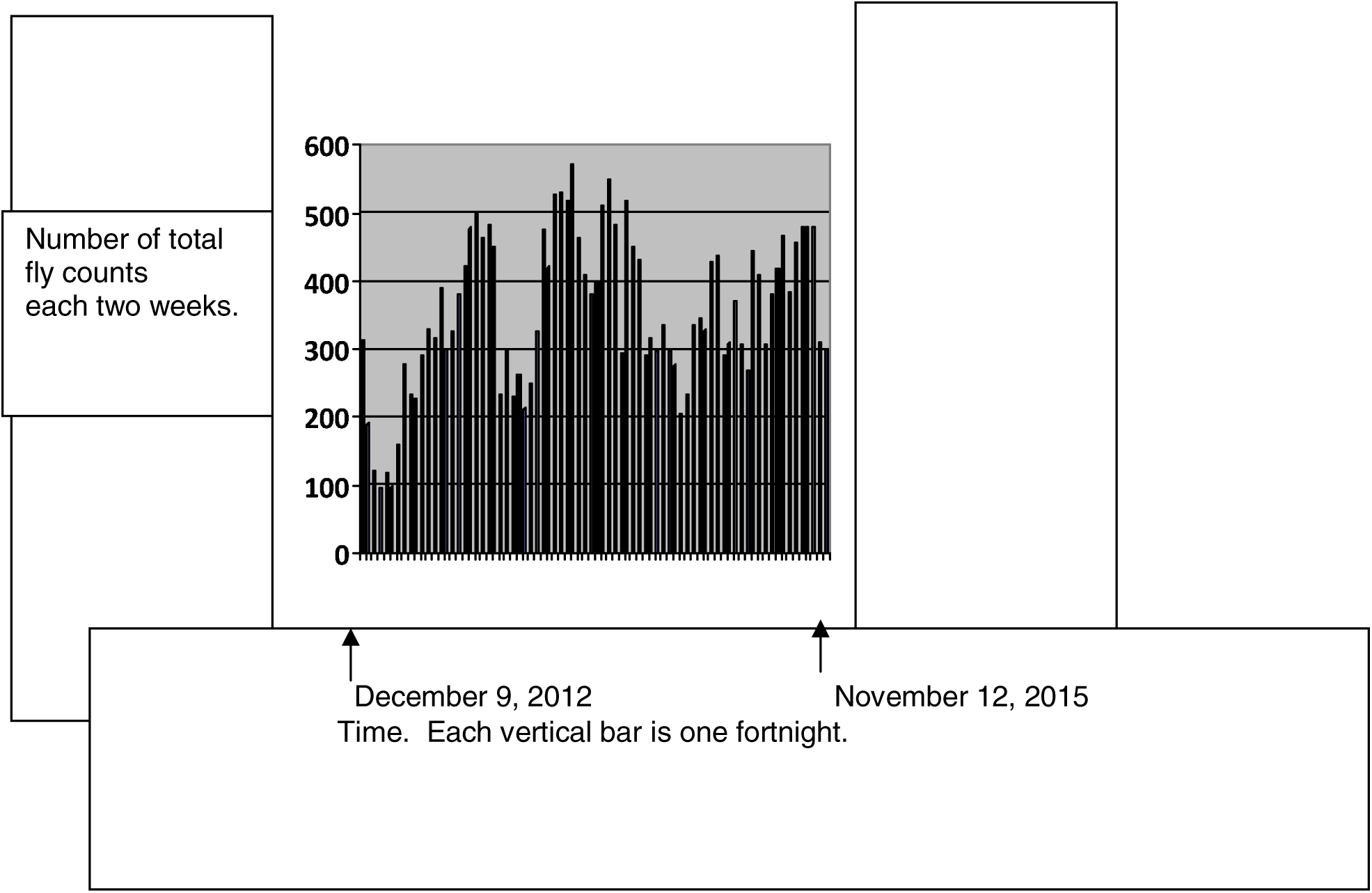
Thirty five months of daily counts of the captive fruit fly population through two windows accumulated at two week intervals. The flies’ food contained decreasing concentrations of a cocktail of chemicals that increased methylation of the fly genome. The noise tends to obscure the overall trend.

By now a poised observer might reasonably remark that all these graphs are beginning to look alike, so let me clarify the data by, once again, lumping observations. This time they are pooled by percentage of the original cocktail offered in the food, beginning with the baseline we saw before. *(fig. 8)*

**Fig. 8.**
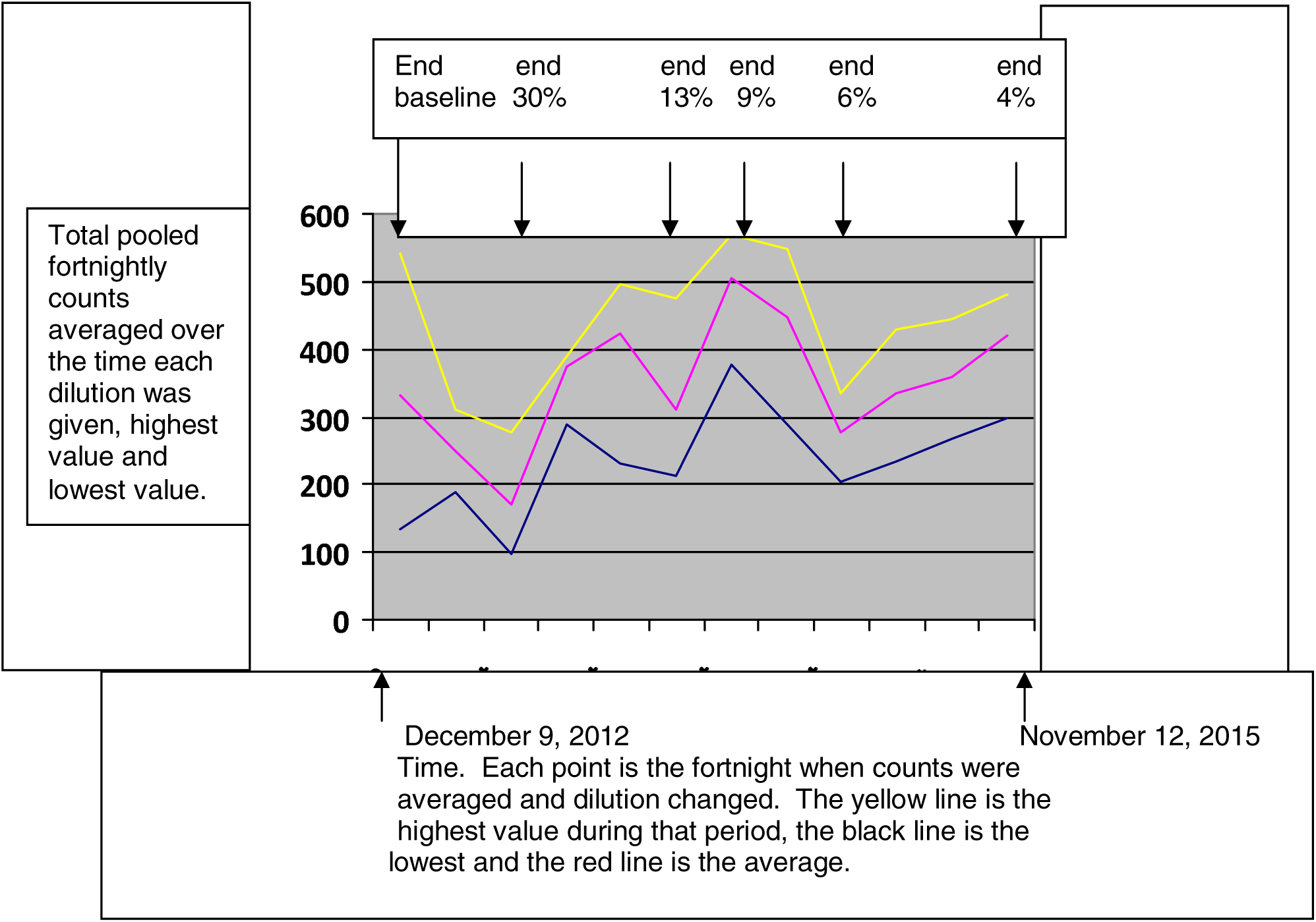
Thirty five months of daily counts of the captive fruit fly population through two windows accumulated at two week intervals, which were averaged over the course of the use of any one dilution with average, highest and lowest values shown. The flies’ food contained decreasing concentrations of a cocktail of chemicals that increased methylation of the fly chromosomes. Selected times when the concentration was reduced are indicated at the date when each concentration was reduced.

Since there is no reason to believe that the data are normally distributed, giving 90% confidence limits would imply that we know more than we actually know, so I have graphed the mean values and highest and lowest values of the fortnightly counts. In fact, except for the baseline, there is very little difference between these and the ninety percent limits. So we shall proceed in chronological order:

“Baseline” is the last day of the baseline counts showing average, highest and lowest fortnights during that time. Since this was more than two years and included an entire cycle of the damped oscillation, we are not surprised that the total range between highest and lowest is large. From then, as we introduce the methylating cocktail there is a rapid fall in population through the 30% dilution. This fall at the outset might be attributed to toxic effect or to the effect that rapid methylation would be expected to produce a rapid increase in mismatch between the methylation patterns in different areas, lowering fertility.

The population more than recovers and then falls again to a valley at the end of 13%. At a stretch this could be some sort of nutritional effect; the methylating cocktail may be providing a boost of some key nutrient. Fertility may be falling as that nutrient is withdrawn. The population then rises through 9%. We have already used up toxic effect and nutrition as possible causes, so we must reject the null hypothesis that methylation has no effect on fertility and accept that methylation is, indeed effective. That established, then we must dismiss toxic effect and nutritional effects as being highly unlikely; it’s all methylation affecting the fertility through the ordinary mechanisms by which kinship determines fertility.

Then the population falls through 6% and rises through 4% . This is now complex behavior; in my hands the computer simulation only permits that when both pre-zygotic and post-zygotic mechanisms are in play. So we have more than we bargained for. The initial question was whether the known effect of kinship on fertility might be due to an epigenetic effect mediated by methylation. The answer is yes, and moreover there are pre-zygotic and post-zygotic elements involved.

Going forward, there are three areas where work could be done: the computer simulation, replication of results and determining the actual location of the methylation sites in the genome.

There is an opportunity for someone who does computer work with relish to improve on the computer simulation above. The most pressing issue is the unstable baseline, which casts a shadow over any results and over reproducibility. The issue might be addressed by diligent search for some combination of parameters that does give a stable baseline, by reducing the stochastic chatter by reducing the number of random variables, by increasing the number of elements involved to improve the statistics or by some combination. This last has been a little problematic since the program as rendered already pushes the limits of the number of sites a 32 digit operating system can address; a 64 digit operating system should be ample if sufficient RAM can be included. Once the baseline issue is laid to rest, it should be possible to enunciate the general principles whereby the results are predictable. And then it should be possible to address the most embarrassing problem with the computer simulation:

The inference of all of this is that in the wild populations are seldom limited by the carrying power of the environment but in general by limited fertility because of factors internal to the individuals. Yet the simulations simulate populations – captive, wild or human – by imposing a maximum population size, effectively a rigid limit to environmental carrying power. This might no longer be a problem once the first problem is fixed. On the other hand, in the present form of the program fertility drops roughly linearly with kinship. That might not be the case in reality. Indeed there is a hint to the contrary; the Sibly curve is after all a curve.

Furthermore a team led by Ann Goodman looked at Swedish data that included what happened when a person became rich. The rich proved quite successful in transferring their socioeconomic states to their descendants, but they had fewer children than a matched population that was not rich (15). This is consistent with the notion that rich people invest more in their children and have fewer children but more grandchildren. As it turned out the number of grandchildren was reduced also. Reasoning from the Sibly curve alone, one might predict that rich people have a broader social horizon than poor, are less likely to marry near kin, and that would account for the difference in the number of children and grandchildren. If we accept that logic and then look at the number of great grandchildren, there is a reduction that is greater than the reduction in children and grandchildren combined. This is little enough to go on, but it is at least a hint that the infertility of reduced kinship increases more than linearly. That could be taken into account with the appropriate computer program.

Replicating the experiment would of course be highly desirable, but problematic as it involved an hour or two a day for years. It would be easier to do the experiment in bottles or vials than in a cumbersome cage. In fact I ran a sort of pilot experiment along side of the cage, with three bottles as the control transferring four males and four females into new bottle each four weeks and three bottles with the same dilutions as the cage, only about a month earlier. (Counts were difficult, so I counted the flies by tens. This is what I found on day 28 during the four months they received a four percent solution. *(table 2)*

**Table 2.**

| Date | I | II | III | IV | V | VI |
| --- | --- | --- | --- | --- | --- | --- |
| August 20, 2015 | 350 | 340 | 270 | 250 | 310 | 360 |
| September 17, 2015 | biofilm | biofilm | biofilm | biofilm | biofilm | biofilm |
| October 15, 2015 | 310 | 290 | 320 | 120 | 100 | 80 |
| November 12, 2015 | 340 | 290 | 420 | 80 | 40 | Dead |
| December 10, 2015 | 230 | 240 | 280 | 200 | 100 | 100 |

Three problems arise. Sometimes bacterial biofilm would invade a bottle. Although frequently the fly population so insulted would survive, presumably because the larvae ate the biofilm, it seemed clear that the bacteria were throwing the counts off, so I exclude them. Once a population in a bottle simply died; this was rare and happened with no biofilm seen. While the fertility in the cage seems to be going up, the fertility in the bottles is going down. I suspect that the flies in the cage adopted a mating strategy different from the one foisted on them by me under the microscope.

A final challenge is to find out at exactly what sites on the DNA the methylation patterns critical to fertility lie. I cannot be of much help, but after years of jousting with the program my impression is that the sites will all be bunched up on one or two chromosomes.

